# Translation control of autophagy genes upon hydroxyurea-induced genotoxic stress

**DOI:** 10.1101/2025.04.09.648074

**Authors:** Gayatri Mohanan, Kumkum Nag, Harsh Singh Senger, Purusharth I Rajyaguru

## Abstract

The fine balance between cellular homeostasis and stress response is crucial for cell survival. In this study, we identify translation regulation of specific autophagy (ATGs) genes by translation repressor Sbp1 upon hydroxyurea-mediated genotoxic stress. Sbp1 localizes to reversible, mRNA-containing granules specifically upon HU stress in RNA recognition motif 1 (RRM1) and RGG motif-dependent manner. Granule localization is independent of eIF4G1 binding despite increased arginine methylation. Deletion of SBP1 increased the tolerance to HU. RNA sequencing of polysome fractions identified ATG1, 2, 9 mRNAs being translationally upregulated in *Δsbp1* upon HU stress. Translation of TEL1 and MEC1, the upstream activators of autophagy, also increased in *Δsbp1*. Concomitantly, increased autophagy and decreased NHEJ repair in *Δsbp1* cells was observed. Together, these findings identify Sbp1 as a regulator of the autophagy pathway in response to genotoxic stress by modulating autophagy at two levels, possibly mediated by its ability to localize to granules.

## Introduction

Cells can experience genotoxic damage from various sources, including environmental factors, chemicals, or even normal cellular processes [1], [2]. The ability of cell to respond to this damage is crucial for maintaining its survival. The response to DNA damage is highly dependent on the type and severity of the insult. In response, specific DNA damage response (DDR) mechanisms are activated, initiating complex signaling pathways that work to either repair the damage or, in more extreme cases, prevent the damaged cell from proliferating [2]. Prolonged exposure to genotoxic stress, could induce cells to enter an adaptive state, adjusting their response mechanisms to manage the ongoing damage and maintain cellular function [3], [4]. One of the first responses to genotoxic stress is cell cycle checkpoint activation mediated by sensor kinases such as Tel1 (ATM) and Mec1 (ATR) in *Saccharomyces cerevisiae* which belong to the PI3K-related protein kinase (PIKK) family of enzymes [5]. Upon activation, these kinases phosphorylate downstream kinases leading to signal amplification and DDR. These enzymes are also reported to directly interact with the repair machinery at the sites of DNA damage to facilitate repair. It has been reported that specific genotoxins can induce autophagy, such as genotoxin-induced targeted autophagy (GTA), mediated via Tel1 and Mec1 [6]. Autophagy is a highly conserved catabolic process that plays a key role in maintaining cellular homeostasis [7], [8]. It recycles macromolecules and even organelles, allowing the broken-down metabolites to be reused. Autophagy is also reported to be involved in stress response and has cytoprotective roles [9], [10]. Autophagy is negatively regulated by Torc1 complex and gets activated upon Torc1 inhibition [11], [12]. Autophagy and DNA damage response are inter-linked. Autophagy is reported to be important for DNA damage response whereas reports also suggest that DNA damage induces autophagy [6], [13], [14]. Several reports indicate that autophagy is critical for homologous recombination mediated double-strand DNA break repair [15], [16]

To understand the cellular response to genotoxic stress, particularly in relation to DNA breaks, various chemicals and drugs are used to induce damage. For instance, hydroxyurea (HU), an inhibitor of ribonucleotide reductase, causes depletion of deoxyribonucleotide triphosphate (dNTP) pools, hindering DNA synthesis and replication fork arrest [17], [18]. Methyl methanesulfonate (MMS), an alkylating agent, methylates DNA, introducing lesions [19]. Cisplatin, a chemotherapy drug, intercalates into DNA and forms platinum-based adducts, disrupting DNA structure and function [20], [21]. Similarly, Zeocin, a member of the bleomycin family of glycopeptides, can intercalate into DNA and induce strand breaks [22]. Camptothecin (CPT), a topoisomerase inhibitor, interferes with the transcription machinery by stalling it on the DNA, which causes a collision with the replication machinery and results in DNA damage[23]. These drugs provide valuable tools for studying the cellular response to genotoxic stress, shedding light on the intricate mechanisms that protect cells from genomic instability. [24], [25], [26][27], [28], [29]

RGG-motif containing proteins are the second largest class of RBPs and are characterized by the presence of RGG/RGX repeats [30]. These are low complexity, intrinsically disordered proteins, a property that allows them to bind a variety of biomolecules such as nucleic acids, proteins and lipids and are therefore involved in various cellular functions [31], [32]. Many of these proteins are also known to localize to cytoplasmic mRNA-protein condensates, such as P-bodies (PBs) and stress granules (SGs). These are higher-order mRNA-protein complexes (mRNPs) that play a crucial role in regulating mRNA metabolism under stress conditions [24], [25], [26]. Upon exposure to various types of stress, the number and size of these condensates significantly increase, and the composition of these granules is reported to be stress-specific [27], [28], [29]. Sbp1 is an RGG-motif containing protein that is known to localize to PBs and SGs upon stresses such as glucose deprivation and sodium azide stress. It is a decapping activator and was recently implicated in PB disassembly [33], [34]. It is also involved in translation repression by binding to eIF4G1, and this activity is modulated by methylation in the RGG motif, but its cellular targets are yet to be determined [35]. In this study, we identify the translation targets of Sbp1 following hydroxyurea (HU)-induced genotoxic stress. Our findings reveal the intricate cellular crosstalk between cytoplasmic mRNA-protein condensates, the DNA damage response (DDR), and autophagy. We explore how these interconnected pathways cooperate to enable the cell to mount an appropriate response to genotoxic stress and adapt accordingly.

## Results

### Sbp1 localizes to granules in response to HU treatment

We tested localization of Sbp1 (endogenously tagged to GFP), in response to several genotoxic stress. Cells expressing Sbp1-GFP were treated with 0.2 M hydroxyurea, 150 µM cisplatin, 0.03% methyl methanesulphonate, 100 µM camptothecin or 100 µg/ml zeocin (as described in Materials and Methods through live cell imaging. We observed localization of Sbp1 to granules upon HU stress, but no changes in localization was observed upon exposure to other genotoxins (Figure 1A and 1B). To measure Sbp1 protein levels in response to different genotoxins, we measured GFP intensity and observed no significant differences (Figure 1C). To determine whether the localization change of Sbp1 in response to HU stress was also observed in other RGG motif-containing proteins, we analyzed the localization of GFP-tagged Gbp2, Npl3, and Psp2 (Figure 1D) under HU treatment, but no changes in their localization were detected (Figure 1E). These findings suggest that the Sbp1 localization change in response to hydroxyurea is both stress- and protein-specific.

**Figure 1:**
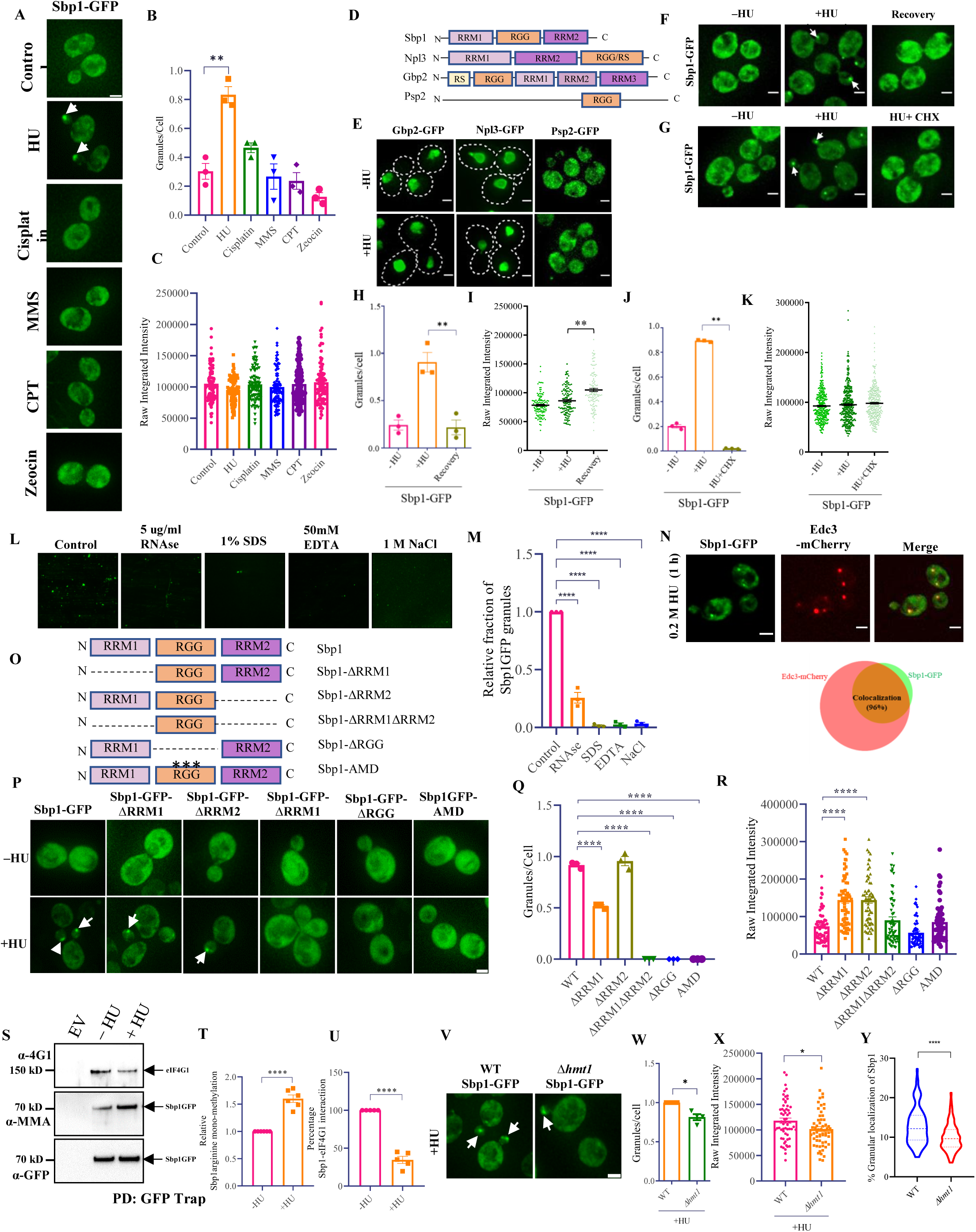
Sbp1 localizes to granules in response to HU induced genotoxic stress. (**A**) Live cell imaging showing localization of GFP tagged endogenous Sbp1, upon treatment with 0.2 M HU, 150 µM cisplatin, 0.03% methyl methanesulphonate (MMS), 100 µM camptothecin and 100 ug/ml zeocin. (**B**) Quantification of granules as granules per cell for Sbp1 foci. (**C**) Quantification of relative GFP fluorescence intensity plotted as raw integrated intensity, from the same experiments. Statistical significance was calculated using Tukey’s multiple comparisons test. Error bars indicate the standard error of mean. n=3; ≥300 cells were counted for quantification of granules per cell and 90 cells were used for GFP fluorescence intensity quantification. (**D**) Schematic representation of domain organization of RGG-motif containing proteins Sbp1, Npl3, Gbp2 and Psp2. (**E**) Live cell imaging showing localization of GFP-tagged Gbp2, Npl3 and Psp2 upon treatment with 0.2 M HU for 120 min. Dotted white line denotes cell boundary. (**F**) Live cell image showing the change in localization of endogenously tagged Sbp1-GFP after HU treatment followed by incubation in non–HU-containing media for 90 min (recovery) (**G**) Live cell image showing the change in localization of endogenously tagged Sbp1-GFP after 5 min of 0.1 mg/ml CHX treatment post 55 min of HU treatment. (**H**) Quantification of granules as granules per cell and (**I**) total GFP intensities upon recovery. (**J**) Quantification of granules as granules per cell (**K**) and total GFP intensities upon CHX treatment. n=3; ≥300 cells were counted for analysis; ≥ 100 cells were used for intensity calculations. (**L**) Sbp1GFP granules enriched from HU stressed cells, treated with 5 µg/ml RNAse, 1% SDS, 50 mM EDTA and 1 M NaCl visualized by fluoresence imaging. (**M**) Quantification of relative GFP puncta in L. (**N**) Live cell image showing co-localization of endogenously tagged Sbp1-GFP with plasmid borne Edc3-mCherry upon 0.2 M HU stress. And schematic representation of Sbp1GFP-Edc3mCherry colocalization. (**O**) Schematic representation of domain organization of WT and domain deletion mutants of Sbp1. (**P**) Live cell image showing the change in localization of plasmid borne WT Sbp1-GFP and domain deletion mutants after 0.2 M HU treatment for 60 min. (**Q**) Quantification of granules as granules per cell and (**R**) total GFP intensities. n=3; ≥300 cells were counted for analysis; ≥ 100 cells were used for intensity calculations. (**S**) Western blot showing immunoprecipitation of GFP-tagged Sbp1 upon HU, probed with anti-monomethylated arginine and anti-eIF4G1 antibodies. (**T**) Quantification of relative Sbp1 mono-methylation upon HU treatment (**U**) Quantification of relative Sbp1-eIF4G1 interaction upon HU stress (n=5) (**V**) Live cell image showing the change in localization of endogenously tagged Sbp1-GFP in WT vs *Δhmt1* strain. (**W**) Quantification of granules per cell, (**X**) total GFP intensities and (**Y**) percentage granular localization of Sbp1GFP (as shown in H) n=4. Error bars indicate standard error of mean and statistical significance was calculated using unpaired t-test. White arrows indicate cytoplasmic foci. Scale bar denotes 2 μm.

To investigate the nature of Sbp1 granules, we tested the reversibility of Sbp1 granule formation by allowing HU-stressed cells to recover in drug-free media. After recovery, we observed a significant reduction in the number of Sbp1 granules, which returned to baseline levels (Figure 1F and 1H). There was a slight increase in the total GFP intensity, suggesting that reduction in granule numbers is not associated with a decrease in protein levels (Figure 1I). We further explored the composition of Sbp1 granules by examining the presence of mRNA in these structures. For this, we used cycloheximide (CHX), which is known to dissociate mRNA-containing dynamic granules [36], [37]. Similar to the recovery process, CHX treatment led to a significant decrease in the number of Sbp1 granules, without affecting the overall protein levels, as indicated by the total GFP intensity (Figure 1G, 1J and 1K). To gain more insight into the nature of Sbp1 granules, we treated Sbp1 granule-enriched fractions *ex vivo* with various reagents, including RNase, SDS, EDTA, and NaCl. RNase treatment leads to RNA degradation and, therefore, dissociates structures that are stabilized by RNA-protein interactions. SDS is a detergent that can disassemble non-aggregate, higher-order structures. Protein aggregates are resistant to 1% SDS treatment and therefore can be differentiated from higher-order protein structures. NaCl and EDTA can disrupt complexes which require electrostatic interactions for stability. Live cell imaging showed a decrease in the number of Sbp1 granules after treatment with RNase, SDS, EDTA, and NaCl, indicating that Sbp1 granules require RNA for stabilization and Sbp1 association with granules is dependent on electrostatic interactions. These observations suggest that Sbp1 assembles into HU-induced higher order mRNA-protein complex (Figure 1L and 1M). These results also indicate that Sbp1 is likely not part of the condensate core because proteins present in the core are, more stably associated and therefore resistant to RNAse, EDTA and NaCl treatment [38]. Considering that P-bodies have been previously reported to form under HU stress [39], we investigated whether Sbp1 granules colocalize with the conserved P-body marker Edc3. Using mCherry-tagged Edc3, we observed 96% co-localization with Sbp1 granules upon HU treatment (Figure 1N), further supporting the idea that Sbp1 granules are closely associated with P-bodies during genotoxic stress.

### Sbp1 localization to granules upon HU stress is dependent on the RGG motif and RRM1 domain

Sbp1 is a modular protein that contains a central RGG motif, flanked by two RNA recognition motif (RRM) domains: RRM1 at the N-terminal and RRM2 at the C-terminal. To investigate which specific domain/motif is responsible for Sbp1 localization to granules in response to HU stress, we generated domain deletion mutants of GFP-tagged Sbp1, expressed under its own promoter (Figure 1O). Live cell imaging indicated that the deletion of the RRM2 domain had no effect on Sbp1 localization to granules, although it did result in elevated protein levels (Figure 1P, 1Q and 1R). In contrast, deletion of the RRM1 domain led to a 50% reduction in Sbp1 granule, despite a noticeable increase in protein levels (Figure 1P, 1Q and 1R). Interestingly, when both RRM1 and RRM2 domains were deleted, or when the RGG motif was removed, Sbp1 localization to granules was completely abolished (Figure 1P, 1Q and 1R). To assess the role of arginine residues in RGG motif which is the site of arginine methylation by Hmt1 [35] we tested arginine methylation-deficient (AMD) mutant wherein 13 arginine residues in the RGG motif are mutated to alanine (see methods for information on the specific arginine residues that are mutated to alanine). This mutant failed to localize to granules (Figure 1P, 1Q and 1R), indicating that the arginine residues and perhaps arginine methylation of the RGG motif are important for Sbp1 localization to granules. Together, these findings demonstrate that both the arginine residues in the RGG motif and RRM1 domain are crucial for the localization of Sbp1 to granules under HU stress.

### Arginine methylation of Sbp1 increases whereas interaction with eIF4G1 decreases upon HU stress

Since the RGG motif is essential for Sbp1 localization to granules and the translation repression activity of Sbp1 is mediated through its binding to the translation initiation factor eIF4G1—an interaction that is regulated by arginine methylation of the RGG motif —we performed immunoprecipitation assay to investigate the effect of HU treatment on Sbp1 methylation and its interaction with eIF4G1. In this assay, we pulled down GFP-tagged Sbp1 and quantified its mono-methylation. Our results from this assay revealed that HU treatment led to a notable increase in the mono-methylation of the arginine residues in the RGG motif (Figure 1S and 1T). This suggests that the methylation status of the RGG motif is modulated in response to HU-induced stress. Additionally, we assessed changes in the interaction between Sbp1 and eIF4G1 under HU treatment. Interestingly, we observed an unexpected decrease in the interaction between Sbp1 and eIF4G1 upon HU treatment, which supports the idea that Sbp1 localization to granules and its function in HU stress is independent of eIF4G1 (Figure 1S and 1U). To further understand the role of Sbp1 methylation on its ability to localize to granules, we checked the localization of Sbp1-GFP in *Δhmt1* strain upon HU stress. Hmt1 is the major methyltransferase in yeast and is known to methylate Sbp1 [35]. We found a modest but significant decrease in Sbp1 localization to granules and Sbp1 protein levels under HU stress in the *Δhmt1* strain (Figure 1V, 1W, 1X and 1Y). These results collectively indicate that arginine methylation within the RGG motif could play a key role in regulating Sbp1 localization to granules during HU stress.

### Deletion of SBP1 increases tolerance to HU stress and de-regulates translation of several mRNAs in response to HU stress

To test if Sbp1 contributes to HU stress response, we conducted a plating assay to quantify the number of colony-forming units (CFUs) of both WT and *Δsbp1* strains upon HU stress. The results revealed a significant increase in the resistance of the *Δsbp1* strain to HU stress compared to the WT strain (Figure 2A). This suggests that the absence of Sbp1 confers tolerance to HU-induced stress. To further confirm this resistance, we performed a growth curve analysis, which showed a significant reduction in the lag time of the *Δsbp1* strain under HU stress (Figure 2B). Upon complementing *Δsbp1* strain with a plasmid-borne SBP1, the lag time was comparable to wild-type yeast strain (Figure 2C), further supporting the idea that Sbp1 plays a role in the cellular response to HU stress. We performed plating assay using methyl methane sulfonate (MMS), a genotoxic agent that does not alter Sbp1 localization to granules. In contrast to the results observed with HU, we did not observe increased tolerance to MMS in *Δsbp1* strains (Figure 2D). This suggests that the role of Sbp1 is specific to HU stress.

**Figure 2:**
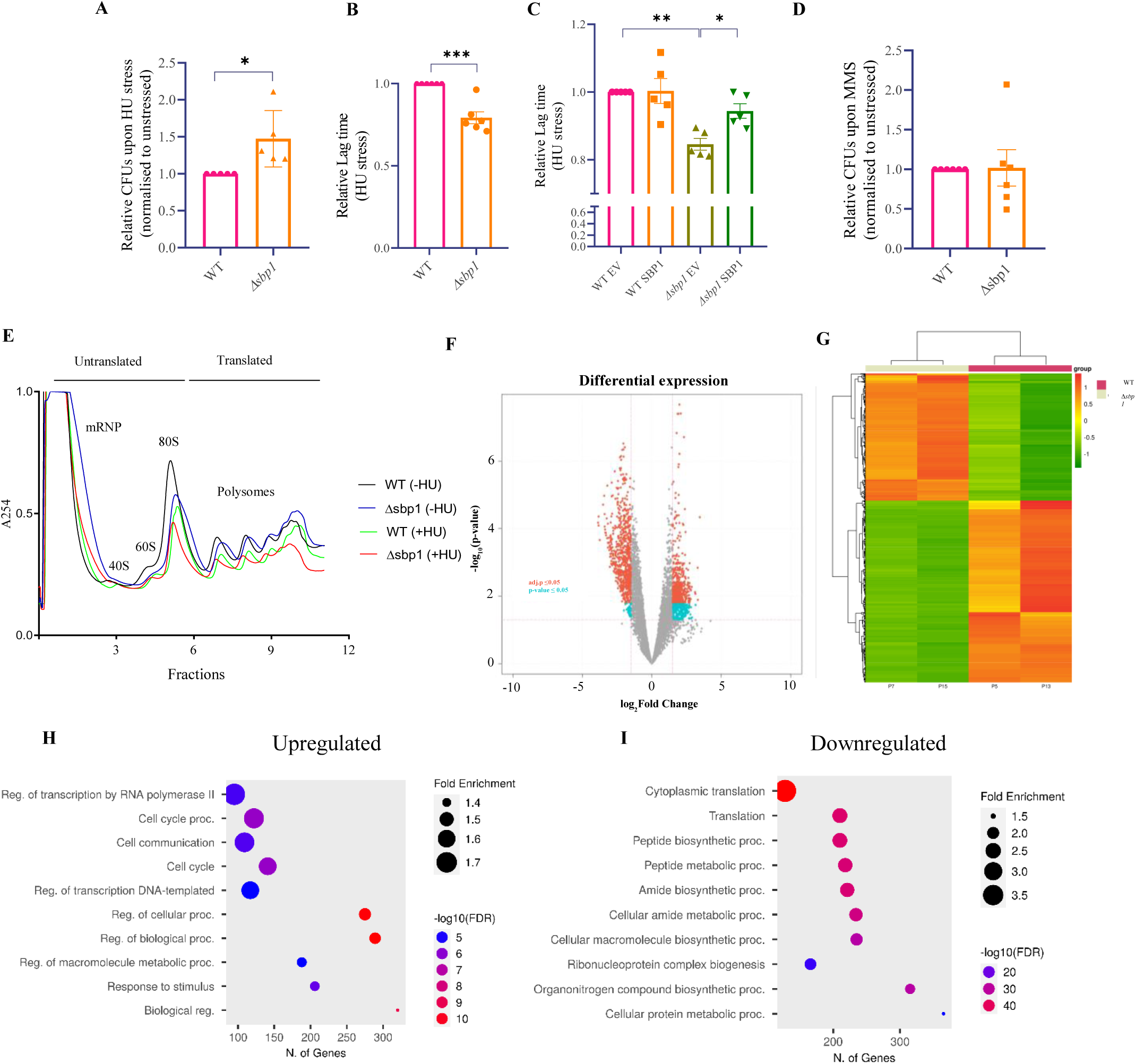
Deletion of SBP1 increases tolerance to HU stress and de-regulates translation of several mRNAs in response to HU stress. (**A**) Quantification of plating assay, plotted as CFUs normalized against WT (n=5). 100 ul of 10^-4^ dilution cell suspension were plated on control (−HU) or 100 mM HU-containing YEPD plates (**B**) Quantification of growth rate in terms of lag time, plotted as change w.r.t. WT (n=6). (**C**) Quantification of growth rate in terms of lag time, upon complementation with plasmid borne-Sbp1, plotted as change w.r.t. WT (n=5). (**D**) Quantification of plating assay, plotted as CFUs normalized against WT (n=5). 100 ul of 10^-4^ dilution cell suspension were plated on control or 0.01% MMS-containing YEPD plates, n=5. (**E**) Representative polysome profiles of WT or *Δsbp1* strain in control or 0.2 M HU treated cells (n=2). (**F**) Volcano plot and (**G**) heat map of mRNAs that are differentially regulated in polysomes upon HU stress in *Δsbp1* strain when compared to WT strain. Bubble plot of gene ontology analysis (biological processes) of (**H**) upregulated and (**I**) downregulated mRNAs. Error bars indicate standard error of mean and statistical significance was calculated using unpaired t-test.

Sbp1 is a translation repressor protein. To investigate, if Sbp1 contributed to HU stress response by modulating the translation of specific mRNAs, we performed polysome profiling followed by RNA sequencing to compare global translation in WT and *Δsbp1* strains upon HU treatment (Figure 2E). Abundance of 1755 protein-coding mRNAs was significantly (with a log2FC ≥ 1) altered in polysome fractions from the *Δsbp1* strain under HU stress. Among these, the abundance of 683 mRNAs was increased, while 1072 mRNAs showed reduced abundance in polysome fractions (Figure 2F and 2G). This was unexpected, as Sbp1 is a translation repressor, and therefore it is anticipated that the number of mRNAs undergoing increased polysome association is likely to be more than those undergoing decreased polysome association. The mRNAs showing increased translation were primarily involved in transcription regulation and cell cycle processes, while mRNA encoding proteins associated with translation were significantly downregulated in the polysome fractions of *Δsbp1* upon HU stress (Figure 2H and 2I).

### SBP1-associated tolerance to HU correlates with increased autophagy and decreased NHEJ repair

Autophagy is reported to enhance cellular growth and life span. Motivated by the observation of increased cellular tolerance in *Δsbp1* cells upon HU stress (Figure 2A and 2B), we analyzed the change in autophagy-associated genes in the translatome data. Five autophagy-related genes were observed to be translationally upregulated in *Δsbp1* upon HU stress. We validated the upregulation of *ATG1, ATG2,* and *ATG9* by RT-qPCR with gene-specific primers (Figure 3A). We observed an average 2 log_2_FC upregulation in the translation of all three mRNAs in *Δsbp1* compared to WT under HU stress (Figure 3A). To ascertain the functional relevance of upregulation, we performed an autophagy assay using a GFP-ATG8 expressing plasmid [40]. Autophagy was monitored in both WT and *Δsbp1* strains expressing this construct by assessing the formation of autophagosomes and the accumulation of free GFP in the vacuole in control and HU-treated conditions. Using live cell imaging, we observed a significant increase in the percentage of cells with vacuolar GFP intensity in *Δsbp1* strains upon HU treatment compared to WT cells (Figure 3B and 3C). Similarly, there was a significant reduction in autophagy induction upon overexpression of SBP1 in both control and HU treated cells (Figure 3D and 3E). These observations, along with mRNA expression analysis in the polysome fractions (Figure 3A) suggest that Sbp1 acts as a negative regulator of autophagy upon HU-induced stress.

**Figure 3:**
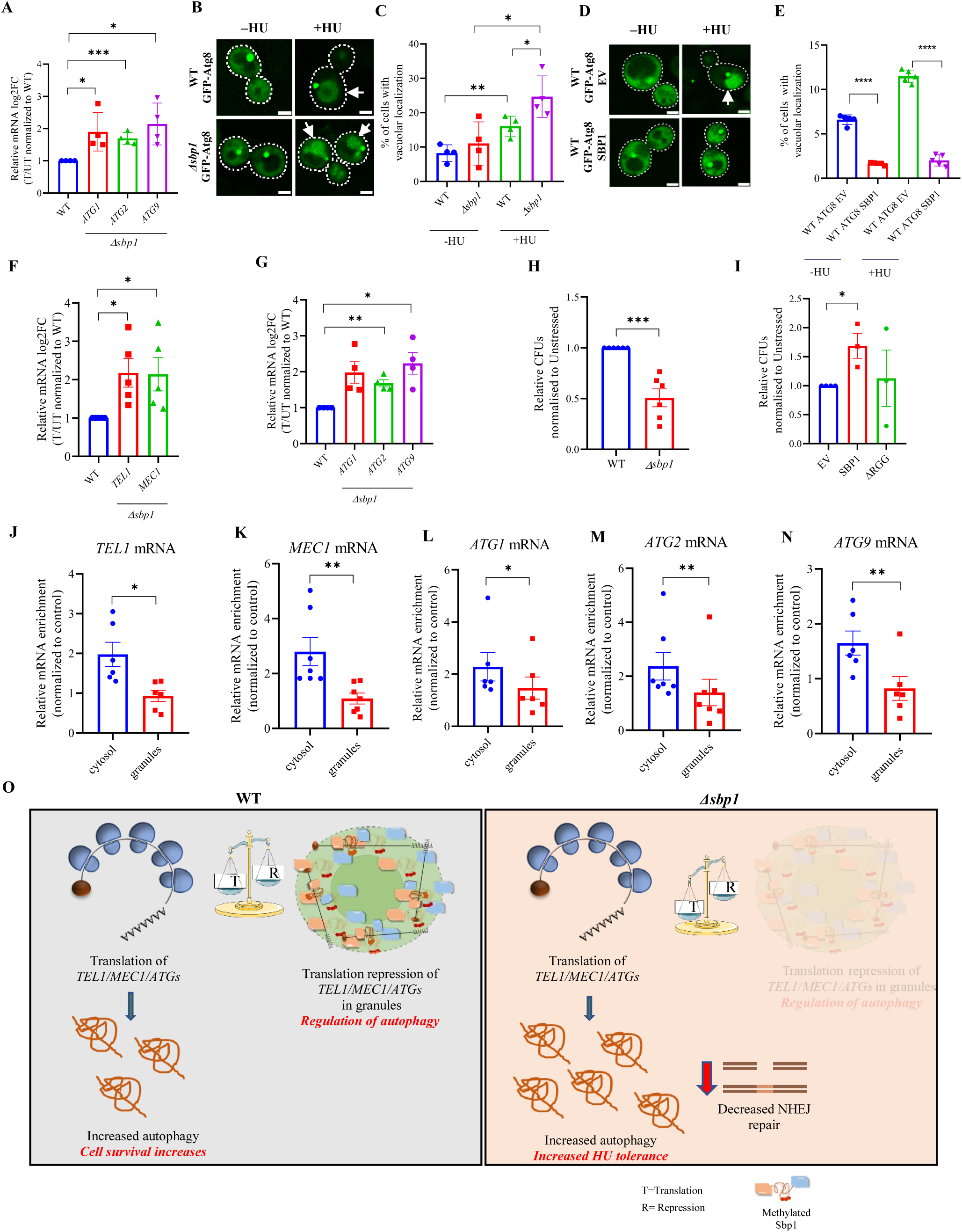
SBP1-associated tolerance to HU correlates with increased autophagy and decreased NHEJ repair: (**A**) Quantification of ATG1, ATG2 and ATG9 mRNAs in the polysome fractions plotted as relative log2-Fold change ratio of Translated/Untranslated fractions normalized to WT, n=4. (**B**) Live cell image showing localization of plasmid borne GFP-ATG8 expressed in either WT or *Δsbp1* strain in control or HU treated cells. (**C**) Quantification of autophagic flux, measured as percentage of cells with vacuolar GFP signal, n=4. (**D**) Live cell image showing localization of plasmid borne GFP-ATG8 expressed in WT strain co-transformed with either EV or plasmid borne SBP1, in control or HU treated cells. (**E**) Quantification of autophagic flux, measured as percentage of cells with vacuolar GFP signal, n=3. (**F**) Quantification of TEL1 and MEC1 mRNAs in the polysome fractions plotted as relative log2-Fold change ratio of Translated/Untranslated fractions normalized to WT, n=5. (**G**) Quantification of ATG1, ATG2 and ATG9 mRNAs from total cell lysate in *Δsbp1* strain normalized to respective controls and to WT, n=4. (**H**) Plasmid recircularization assay, quantified as relative CFUs normalized to respective controls and to WT, n=6. (**I**) Quantification of NHEJ activity in suicide deletion strains expressing either EV, SBP1 or RGG motif deletion mutant, n=3. (**J-N**) Quantification of relative mRNA enrichment of TEL1 (n=7), MEC1 (n=7), ATG1 (n=6), ATG2 (n=7) and ATG9 (n=7) mRNA in cytosolic and granule enriched fraction, normalized to unstressed control. Error bars indicate standard error of mean and statistical significance was calculated using unpaired t-test. (**O**) Schematic representation of Sbp1 mediated regulation of autophagy and its possible role in cellular tolerance to HU.

HU treatment is reported to induce autophagy [6], [41]. DNA damage response triggers a specific type of autophagy known as genotoxin-induced targeted autophagy (GTA), which is mediated by checkpoint kinases like Tel1, Mec1, and Rad53 leading to transcriptional upregulation of ATGs [6]. To determine if *Δsbp1*-induced tolerance to HU could be linked to GTA, we first analyzed and validated the upregulation of Tel1 and Mec1 by RT-qPCR with gene-specific primers. We observed an average log_2_FC=2 upregulation in the translation of both *TEL1* and *MEC1* mRNA in *Δsbp1* compared to WT under HU stress (Figure 3F). To test if the increase in TEL1 and MEC1 expression led to increased transcription of ATGs, we isolated total RNA from WT and *Δsbp1* strains and quantified the expression of *ATG1*, *ATG2* and *ATG9* mRNAs. As expected, there was a significant increase in the expression of these mRNAs in *Δsbp1* strains compared to WT upon HU stress (Figure 3G). Together, our results indicate that increase in autophagy in *Δsbp1* upon HU stress is also correlated with increased translation of *TEL1* and *MEC1* mRNA followed by increased expression of ATGs pointing towards multi-level regulation of autophagy by Sbp1. Since autophagy and DNA repair are interlinked, and we observe HU tolerance in *Δsbp1* compared to WT, we assessed the non-homologous end joining (NHEJ) mediated DNA repair in *Δsbp1* compared to WT. Strains were transformed with enzymatically linearized plasmid and plated on selection media. The number of colony forming units (CFUs) that appeared on the plate, which is proportional to the extent of repair activity, was quantified. We observed a significant reduction in NHEJ activity upon HU stress (Figure 3H). Similarly, using a suicide deletion strain which scores for NHEJ mediated repair, we observed a significant increase in NHEJ activity upon exogenous expression of plasmid borne SBP1 (Figure 3I). This increase in NHEJ activity was not observed when RGG-motif deletion mutant, which also fails to localize to granules, was expressed (Figure 3I). These observations indicate a reciprocal regulation of autophagy and NHEJ mediated repair. Since it is reported that autophagy augments homologous recombination (HR)[15], we believe that decreased NHEJ activity in *Δsbp1* strain upon HU treatment is a result of increased HR activity mediated by upregulation of HR repair pathway. Moreover, TEL1 and MEC1, which are also upregulated in *Δsbp1* strain upon HU, could also augment HR mediated DNA repair. Increased HR repair, which is known to be the error-free repair, could also lead to HU tolerance associated with *Δsbp1*.

To understand the role of Sbp1 localization to granules and regulation of autophagy, we performed granule enrichment in WT cells upon HU treatment and scored for partitioning of *TEL1*, *MEC1*, and *ATG1/2/9* mRNAs to cytoplasm and granular fraction. Granule enrichment was confirmed by measuring the levels of Sbp1 in cytosol vs granule enriched fraction (data not shown). P-bodies are known to be sites of mRNA storage and decay, and cytoplasm is usually associated with translation. Our results indicated a differential partitioning of *TEL1*, *MEC1*, and *ATG1/2/9* mRNAs to the cytoplasm, but a significant part of the mRNA pool was present in the granule enriched fraction as well (Figure 3J-3N). This indicates that a larger fraction of these mRNAs undergoes active translation upon HU stress which is in line with existing reports which show autophagy induction. But our results indicate that a part of these mRNA pools are stored in the P-bodies in an inactive form. Upon deletion of Sbp1, more mRNAs are able to translate and therefore we observe a significant increase in the translation of these genes (Figure 3O). Together, these results highlight the intricate relationship between autophagy, DNA repair and P-bodies.

## Discussion

The results from this study reveal critical insights into the role of Sbp1 in regulating cellular responses to genotoxic stress, particularly in response to hydroxyurea (HU)-induced DNA damage. We have observed that Sbp1 localizes to cytoplasmic granules specifically under HU stress, which was not observed with other genotoxic treatments such as cisplatin, methyl methanesulfonate, camptothecin, or zeocin (Figure1A). This specificity highlights the selective response of Sbp1 to HU-induced stress, providing an important clue into its function during DNA damage. Importantly, we observed that the granule formation was reversible upon recovery in drug-free media and contained mRNA (Figure 1F and 1G), indicating the dynamic nature of these granules. Additionally, the treatment of granule-enriched fractions with RNase, EDTA and NaCl revealed that Sbp1 is associated with the outer shell of granules under HU stress, and not part of the core granule complex, which is typically more resistant to dissociation (Figure 1L and 1M). This observation is also backed by the fact that Sbp1 has a relatively lower partition coefficient and very high recovery dynamics [42]. This property probably enables the transition of Sbp1 and its bound mRNA targets between granules and cytoplasmic mRNA pool (discussed later). Our co-localization experiments with a conserved P-body marker Edc3 indicates that Sbp1 localizes to P bodies upon HU stress (Figure 1N). Moreover, our results suggest that arginine methylation is crucial for granule formation during HU stress, a mechanism that is independent of Sbp1-eIF4G1 interaction, a translation initiation factor (Figure 1T and 1U). This is an interesting and somewhat unexpected finding, as previous studies have implicated Sbp1 interaction with eIF4G1 in its role as a translation repressor. Our data, however, indicate that the function of Sbp1 in HU stress is independent of this interaction, suggesting a distinct regulatory mechanism for Sbp1 in response to genotoxic stress, which might also play a role in condition-dependent target specificity.

The deletion of SBP1 was found to significantly increase tolerance to HU stress, as evidenced by increased colony-forming ability and a reduced lag time in growth curves (Figure 2A and 2B). Interestingly, this increased resistance was specific to HU stress, as no enhanced tolerance was observed with methyl methanesulfonate (MMS), a genotoxic agent that does not alter Sbp1 localization to granules (Figure 2D). This specificity underscores the targeted role of Sbp1 in modulating the response to HU stress by affecting the translation of specific mRNAs. We identify *ATG1, ATG2* and *ATG9* as specific translation targets of Sbp1 (Figure 3A). ATG1 is reported to be involved in the activation of genotoxin-induced targeted autophagy (GTA), a process that helps cells manage DNA damage by degrading damaged proteins and organelles [6]. The autophagy flux assay provides functional validation of increase in autophagy activity in *Δsbp1* (Figure 3B) whereas overexpression of SBP1 had the opposite effect (Figure 3D). Translational upregulation of Mec1 and Tel1 kinases (Figure 3F), which are implicated in GTA and are activators of autophagy genes, further confirms the role of Sbp1 modulating autophagy upon HU stress. Transcriptional upregulation of autophagy genes by Tel1 and Mec1 reported earlier is also captured in the *Δsbp1* background (Figure 3G), which highlights that Sbp1 modulation affects autophagy at the level of both transcription and translation in response to HU. Our investigation also uncovers the reciprocal regulation of NHEJ and autophagy (Figure 3H and 3I). It is reported that HR and NHEJ can compensate for each other’s function in a condition specific manner [43], [44]. Although, in this report, it is not clear if down regulation of NHEJ leads to increased HR, but is speculated. Finally, we demonstrate that the mRNA targets of Sbp1 partition to both cytoplasm and granules which could enable Sbp1 and its bound targets to transit between translationally active and inactive states. This speculation is further supported by the ability of Sbp1 to loosely associate with P-bodies (Figure 1L). Although, the mechanistic details of this movement need further investigation.

We currently do not understand the physiological relevance of Sbp1 mediated down-regulation of autophagy in response to HU stress. We speculate that Sbp1 may act as a molecular brake on autophagy under genotoxic stress, preventing its overactivation and ensuring that the cell can adequately manage DNA damage without triggering excessive degradation of cellular components and maintain cellular homeostasis under stress. Further investigation is required to identify the precise role of Sbp1 localization to granules and to identify the mechanism of Sbp1 mediated translation repression of ATGs upon HU stress. Even though we demonstrate the correlation between NHEJ mediated repair and Sbp1, the molecular mechanism of this interaction needs further investigation. Another limitation of this study is that it lacks the exact role of Sbp1 granules in regulation of autophagy. Currently, we do not explore the mRNA and protein components of Sbp1 granules and how the composition of these granules can affect targets of Sbp1. Moreover, identification of other protein like Sbp1, in higher order, complex systems such as mammalian models which could be involved in similar regulatory pathways remains an important future direction.

Overall our results identify a new example of autophagy pathway gene regulation at the translation and transcriptional level. We establish several ATG genes (such as ATG1, 2 and 9) as translation targets of Sbp1 in HU stress. Nucleolin is a structural homolog of Sbp1, and therefore, addressing the role of this protein in translation control of autophagy genes would be an obvious starting point. As the field continues to explore the molecular players involved in regulation of autophagy and DNA repair, our work sets the stage for further investigations into the translational control of these pathways, potentially offering therapeutic avenues for diseases involving mis-regulated stress responses.

## Methods and Materials

### Yeast transformation

All the strains and plasmids used in the study are listed in supplementary Table T1 and T2, respectively. The strains were cultured to an OD600 of 0.6 in yeast extract and peptone with glucose (YEPD), then pelleted. The cells were resuspended in 100 mM Lithium Acetate and divided into 50 µL aliquots. To each aliquot, 240 µL of 50% PEG (v/v), 36 µL of 1 M LiAc, 25 µL of salmon sperm DNA (100 mg/mL), and 100 ng of the respective plasmid DNA were added, and the mixture was vortexed. The cells were incubated at 30 °C for 30 min, followed by 15 min at 42 °C. After incubation, the cells were pelleted, resuspended in 100 µL of water, and plated on synthetic defined media with glucose agar plates. The plates were incubated at 30 °C for 2 days until colonies appeared.

### Drug treatments

Yeast cultures were grown in 10 mL of SD-Leu-media containing glucose, to an OD600 of 0.6-0.8, and then divided into two equal portions. Each portion was treated with 0.03% MMS, 0.2 M HU, 150 µM cisplatin, or 150 µM zeocin, along with a vehicle control, and incubated for 60 min at 30 °C. For CHX treatment, after incubation with HU, 3 µL of a 100 mg/mL cycloheximide solution (dissolved in methanol) or methanol (vehicle control) was added to the HU-treated fractions. The cells were incubated on a shaker for 5 min, then pelleted for live-cell imaging.

### Yeast live cell imaging

After the incubation period, yeast cells were harvested by centrifugation at 18,000 g for 15 seconds, resuspended in 20 µL of the remaining media, and spotted onto a glass coverslip (No.1) for live-cell imaging. Images were captured using a Deltavision RT microscope system with SoftWoRx 3.5.1 software (Applied Precision, LLC), utilizing an Olympus 100× oil immersion objective with a 1.4 NA. The Green Fluorescent Protein (GFP) channel was set to exposure times of 0.2 or 0.6 seconds, with transmittance set at 32% or 50%. A minimum of 80-100 cells were analyzed for each experiment. Granules per cell were quantified, and the total GFP fluorescence intensity in the cells were determined using ImageJ software. Statistical analysis was performed with GraphPad Prism Version 7.0.

### Western blotting

SDS-polyacrylamide gels were run and transferred onto Immobilon-P Transfer Membranes® (MERCK) using a Transfer-Blot® Semi-Dry Transfer System (BIO-RAD). The transfer process was performed at 10 V for 1 hour to ensure efficient protein transfer. After transfer, the membrane was blocked with 5% skimmed milk to prevent non-specific binding. Following the blocking step, the membrane was washed to remove any excess blocking solution and then incubated with the primary antibody specific to the target protein. In cases where the blot needed to be probed with multiple antibodies, the membrane was stripped of any bound antibodies, re-blocked with 5% skimmed milk, and then re-incubated with the additional antibodies for detection.

### Sbp1-GFP Pull down

Yeast strain expressing GFP-tagged endogenous Sbp1 or WT control were grown in 100 mL of yeast extract-peptone (YEP) media until they reached an OD_600_ of 0.8. The cultures were then divided into two equal parts: one was treated with 0.2 M hydroxyurea (HU) to induce replication stress, while the other was left untreated as a control. After treatment, the cells were harvested by centrifugation, and lysed in a lysis buffer [10 mM Tris, 150 mM NaCl, 1X EDTA-free protease inhibitor complex, RiboLock RNase inhibitor (Cat. No. EO0381)], using bead beating for cell disruption. The lysate was then centrifuged at 2800 g for 5 min. The supernatant was transferred to a fresh microcentrifuge tube, diluted 1:1 with lysis buffer, and 10 µL of GFP-TRAP magnetic agarose beads (Cat. No. GTMA) were added to a final volume of 1 mL. The samples were nutated at 4°C for 120 min, allowing binding to the beads. The beads were then separated using a magnetic stand and washed twice with lysis buffer. Finally, the beads were resuspended in 120 µL of lysis buffer for further analysis.

### Growth curve and plating assay

For the growth curve analysis, secondary yeast cultures were initially grown to the mid-log phase (OD600 between 0.4 and 0.6) in YEPD (Yeast Extract Peptone Dextrose) medium or in synthetic defined drop-out media for complementation growth curves. Once the cultures reached the mid-log phase, they were sub-cultured into fresh media to achieve a final optical density of 0.1 OD600. The cultures were then divided into two equal portions: one was treated with 0.2 M hydroxyurea (HU) to reach a final concentration of 0.2 M, while the other was treated with water as a vehicle control. 200 µL aliquots from each condition were transferred into 96-well clear plates (Corning® Costar Clear Polystyrene 96-Well Plates, Cat ID: 3370). These plates were incubated in a plate reader (Tecan Infinite 200 PRO) at 30°C, with orbital shaking (4 mm) for 40 seconds every hour to maintain a consistent environment. OD600 readings were recorded every hour for a total duration of 24 hours to track the growth of the yeast cultures under both HU treatment and control conditions. The collected data were then analyzed and graphed using GraphPad Prism 9 to determine the growth rates and compare the effects of hydroxyurea treatment against the control.

### Polysome fractionation

Wild-type (WT) or *Δsbp1* strains were grown to an OD600 of 0.8 and treated with 200 mM hydroxyurea (HU) for 60 min. Afterward, the cells were treated with 0.1 mg/mL cycloheximide for 30 min. The cells were then lysed in lysis buffer containing 10 mM Tris (pH 7.4), 100 mM NaCl, 30 mM MgCl_2_, RNase inhibitor, cycloheximide, and an EDTA-free protease inhibitor complex. The pre-cleared lysate was loaded onto a 10–50% sucrose density gradient, followed by centrifugation in an SW41 rotor (Beckman-Coulter) at 260343 g for 2 hours at 4°C. After centrifugation, the gradients were fractionated into two main groups: untranslated fractions (up to the 80S region) and translated polysome fractions. These fractions were collected and pooled, and RNA was isolated from each using the TRIzol-chloroform method. Total RNA was also isolated from the lysate as a reference. RNA samples were treated with DNase to remove any genomic DNA, followed by DNA library preparation using random primers. RT-qPCR was then performed with gene-specific primers to assess mRNA levels in the polysome fractions. The mRNA enrichment in the translated (polysome) fraction was calculated relative to the untranslated fraction. ΔCt values were calculated by subtracting the Ct value of the PGK1 primer (used as an internal control) from the Ct value of the target gene primer. ΔΔCt values were determined by subtracting the ΔCt of total RNA from the ΔCt of the polysome fractions (both translated and untranslated). The final mRNA enrichment values were calculated by normalizing the 2^(-ΔΔCt) ratio of the translated fraction to the untranslated fraction

### RNA isolation, cDNA library preparation and RT-qPCR

RNA was isolated using the TRIzol reagent (G Biosciences, Cat ID 786-652) in combination with chloroform, following a standard extraction protocol. Briefly, the lysate or polysome fractions were mixed with 1 mL of TRIzol reagent and 200 µL of chloroform. The mixture was vortexed thoroughly, followed by flash freezing and then centrifuged at 18,000 g for 20 min to separate the aqueous and organic phases. RNA was precipitated from the aqueous phase by adding isopropanol, followed by flash freezing to facilitate the precipitation process. The samples were then centrifuged at 18,000 g for 30 min at 4°C to pellet the RNA, after which the pellet was washed once with 70% ethanol. The RNA pellet was air-dried and resuspended in an appropriate volume of DEPC-treated water. The RNA was treated with DNase (Thermo Scientific, Cat ID: EN0521) according to the manufacturer’s instructions. Following DNase treatment, equal volume of DNase-free RNA was used for cDNA synthesis. The cDNA libraries were prepared using random primers to specifically amplify mRNA. Once the cDNA libraries were prepared, they were diluted as follows: 1:5 for lysate samples and 1:1 for polysome fractions. These diluted libraries were then used as templates for quantitative reverse transcription PCR (RT-qPCR) in a 10 µL reaction volume. RT-qPCR was conducted using TB Green Premix Ex Taq II (Tli RNase H Plus) (TaKaRa, Cat ID RR820B) for 40 cycles, following the manufacturer’s protocol. Gene-specific primers (Bioserve, India) were used to selectively amplify the target mRNA sequences. The sequences of all the oligonucleotides used in this study are listed in supplementary Table T3. The RT-qPCR data were then analyzed to quantify mRNA distribution across different fractions. Quantification of mRNA enrichment in the polysome fractions (translated fraction) was calculated w.r.t to mRNA in untranslated fractions. ΔCt values for each gene tested were calculated by subtracting Ct value of the PGK1 primer (internal control) from the test primer. The final values were then obtained by normalizing the 2^^(-ΔCt)^ of translated by untranslated polysome fractions and plotted after normalizing with WT.

### RNA sequencing and bioinformatic analysis

The RNA quality assessment was done using RNA ScreenTape System (Catalog: 5067-5576, Agilent) in a 4150 TapeStation System (Catalog: G2992AA, Agilent). 1µl of RNA sample was mixed with 5µls of RNA ScreenTape Sample buffer (Catalog: 5067-5577) and heat denatured at 72 ^0^C for 3 min and immediately placed on ice for 2 min and loaded the sample on the Agilent 4150 TapeStation instrument. The integrity of RNA is determined by RNA integrity number (RINe) assigned by the software. The sequence data was generated using Illumina Novoseq. Data quality was checked using FastQC and MultiQC software. The data was checked for base call quality distribution, % bases above Q20, Q30, %GC, and sequencing adapter contamination. Raw sequence reads were processed to remove adapter sequences and low-quality bases using fastp. The QC passed reads were mapped onto indexed Saccharomyces cerevisiae S288C reference genome (https://www.ncbi.nlm.nih.gov/genome/15? genome_assembly_id=22535) using STAR (Version 2) aligner. On average 97.40% of the reads aligned onto the reference genome. Gene level expression values were obtained as read counts using feature-counts software. For differential expression analysis the biological replicates were grouped as Reference and Test. Differential expression analysis was carried out using the edgeR package after normalizing the data based on trimmed mean of M (TMM) values. Genes with adjusted p-value ≤ 0.05 and absolute log2 fold change ≥ 1 were tested.

### GFP-ATG8 autophagy assay

The WT or *Δsbp1* strains were transformed with a GFP-ATG8 expressing plasmid. The transformants were cultured in synthetic uracil dropout media (SDU^-^) and grown to an OD_600_ of 0.8 from an initial inoculum. Similar methodology was used to measure autophagy upon SBP1 overexpression. WT strain was transformed either with EV or SBP1 on a plasmid under its own promoter, along with GFP-ATG8 plasmid. The transformants were cultured in synthetic uracil-leucine dropout media (SDUL^-^) and grown to an OD_600._ Once the cultures reached the desired density, they were divided into three groups: one was treated with 0.2 M HU for 90 min, another was treated with 200 ng/ml rapamycin (autophagy induction control), and the third served as a control with no treatment. After drug treatment, 1 ml of cells were pelleted down, resuspended in 20 µl media and spotted on a glass coverslip for imaging. Extent of autophagy was determined by calculating the percentage of cells with vacuolar GFP signal.

### Plasmid recircularization assay

1 µg of pRS316 plasmid linearized by NcoI restriction enzyme (as per manufacturers protocol) at *URA3* locus, or uncut plasmid was transformed in equal number of WT or *Δsbp1* cells and plated on Ura drop-out synthetic media in the presence or absence of 100 mM HU. The plates were incubated at 30 °C for 2 days until colonies appeared. The colonies were counted and plotted as a ratio of cut by uncut colonies.

### NHEJ suicide deletion assay

The assay was adapted from Karathanasis et al., 2002 [45]. In brief, the suicide deletion strain YW714 (kindly provided by Professor Thomas Wilson) was transformed with either an empty vector or Sbp1-GFP expressed from a CEN plasmid. Stationary phase cells were plated onto synthetically defined agar plates lacking leucine and adenine (leu- ade+, leu- ade-, or leu- ade-dropout plates). These plates were supplemented with 100 mM hydroxyurea (HU) and one of two carbon source combinations: 2% sucrose, or 1% sucrose combined with 1% galactose to induce DNA double-strand breaks. The plates were incubated at 30°C for 3-4 days, until colonies were visible. Colony counts were taken from all plates and plotted as the ratio of CFUs from HU-treated versus untreated samples, with the values normalized to those obtained from the empty vector control.

### Granule enrichment

The granule enrichment protocol was adapted from Wheeler *et al* [46].WT strain expressing GFP tagged Sbp1 was grown to an OD_600_ of 0.8 from an initial inoculum. The cultures were split into two parts: one without treatment and the other part treated with 0.2 M HU. Post treatment cells were lysed in lysis buffer [10 mM Tris, 150 mM NaCl, 1X EDTA-free protease inhibitor complex, RiboLock RNase inhibitor (Cat. No. EO0381)] using bead-beating and spun at 2000g for 2 min followed by another high-speed spin at 18000g for 10 min. The supernatant was collected and labelled as cytosolic fraction. The pellet was washed with lysis buffer again at 18000g for 10 min. The final pellet was resuspended in 100 µl lysis buffer. A small aliquot of the fractions was also used for SDS-PAGE followed by western blotting to confirm granule enrichment. RNA was isolated from whole cell lysate, cytosolic and granule enriched fraction as described earlier (TRIzol method).

## Supporting information

Supplemental Table S1 (List of strains)

Supplemental Table S2 (List of plasmids)

Supplemental Table S3 (List of primers)

## Acknowledgement and Funding

The authors thank the Rajyaguru lab for their critical inputs and suggestions during this work. This work was supported by the CEFIPRA grant to PIR (7003-H/7003-2); India Alliance DBT-Wellcome trust (IA/I/12/2/500625) and BT/PR51975/BMS/85/23/2024 from the Department of Biotechnology (DBT), Government of India. The authors acknowledge DST-FIST program to the Department of Biochemistry, Indian Institute of Science. GM and HS were supported by a fellowship from MHRD. KN was supported by DBT, India.

## Data availability

The processed sequencing dataset generated in this study has been deposited in Biostudies with the accession number S-BSST1938 (doi:10.6019/S-BSST1938)

## Abbreviations

DDR: DNA Damage response
GTA: Genotoxin associated targeted autophagy
HU: Hydroxyurea
mRNPs: mRNA protein complexes
NHEJ: Non-homologous end joining
P-bodies: Processing bodies

