## Supplemental Table S1 (List of strains) for "Translation control of autophagy genes upon hydroxyurea-induced genotoxic stress"

**Table T1: Strains used in this study**

| **Name** | **Genotype** | **Source** |
| --- | --- | --- |
| yPIR1 | *MATa his3Δ1 leu2Δ0 met15Δ0 ura3Δ0(‘BY4741’)* | Poornima *et al*, 2016 |
| yPIR171 | *MATa his3Δ1 leu2Δ0 met15Δ0 ura3Δ0 hmt1∆::KanMX* | Poornima *et al*, 2016 |
| YW714 | *MATa ade2::SD2-::URA3 his3Δ1 leu2Δ0 ura3Δ0* | Karathanasis *et. al.,* 2002 |
| YW713 | *YW714 MAT yku70Δ::kanMX4* | Karathanasis *et. al.,* 2002 |
| yPIR9 | *MATa his3Δ1 leu2Δ0 met15Δ0 ura3Δ0:*SBP1-GFP (HIS) | Bhatter *et a*l, 2019 |
| yPIR137 | *MATa his3Δ1 leu2Δ0 met15Δ0 ura3Δ0:*PSP2-GFP (HIS) | This study |
| yPIR10 | *MATa his3Δ1 leu2Δ0 met15Δ0 ura3Δ0* SBP1-GFP (HIS) hmt1∆::KanMX | Bhatter *et a*l, 2019 |
| yPIR25 | *MATa his3Δ1 leu2Δ0 met15Δ0 ura3Δ0 sbp1∆*::KanMX | Roy *et a*l, 2022 |
