## Supplemental Table S2 (List of plasmids) for "Translation control of autophagy genes upon hydroxyurea-induced genotoxic stress"

Table T2: List of plasmids used in the study

| **Plasmid**  **number** | **Description** | **Vector** | **Source** |
| --- | --- | --- | --- |
| pPIR92 | CEN plasmid used as empty vector. *URA3*; Ampicillin resistance gene | pRS316 | This study |
| pPIR22 | Sbp1-GFP; *LEU* selection marker; AmpR | pRS315 | Bhatter et al, 2019 |
| pPIR20 | Empty vector; AmpR | pPRS315 | Garg et al, 2022 |
| pPIR24 | Sbp1 ORF; *LEU* selection marker; AmpR | pPRS315 | Bhatter et al, 2019 |
| pPIR52 | Edc3-mCherry; *URA* selection marker; AmpR | pRS416 | Garg et al, 2022 |
| pPIR67 | Sbp1-GFP∆RRM1; *LEU* selection marker; AmpR | pRS315 | This study |
| pPIR68 | Sbp1-GFP∆RRM2; *LEU*selection marker; AmpR | pRS315 | This study |
| pPIR 220 | Sbp1-GFP∆RRM1+2; *LEU* selection marker; AmpR | pRS315 | This study |
| pPIR22 | Sbp1-GFP∆RGG; *LEU* selection marker; AmpR | pRS315 | Bhatter et al, 2019 |
| pPIR23 | Sbp1-GFP AMD; *LEU* selection marker; AmpR | pRS315 | Bhatter et al, 2019 |
| pPIR120 | GFP-Npl3, *URA* selection marker; AmpR | pPS1372 | Kind gift from Hans Krebber Lab |
| pPIR120 | GFP-Gbp2, *URA* selection marker; AmpR | pPS1372 | Kind gift from Hans Krebber Lab |
