## Supplemental Table S3 (List of primers) for "Translation control of autophagy genes upon hydroxyurea-induced genotoxic stress"

**Table T3: List of primers used in this study**

| **Primer** | **Sequence** | **Use** | **Source** |
| --- | --- | --- | --- |
| PIR-KN-Atg1_S | GTCTGTCGGTACAGTGGTATTC | RT-qPCR | This study |
| PIR-KN-Atg1-AS | CGTTATGACATCGTTTGCTCTTT |  |  |
| PIR-KN-Atg2_S | AAGGGTGGCAATGTCTTCTT | RT-qPCR | This study |
| PIR-KN-Atg2_AS | CGAATAGTTCCTCGGTGTTCTC |  |  |
| PIR-KN-Atg9_S | GATGGAGACCACCAGCTAAAT | RT-qPCR | This study |
| PIR-KN-Atg9_AS | GGCACAGAATCATCGGTAGAG |  |  |
| PIR-KN-Mec1_S | CGGAGAAAGCAGACAGAAAGA | RT-qPCR | This study |
| PIR-KN-Mec1_AS | CTCGCATAGGTCCTTGTCTAAAT |  |  |
| PIR-GM-Tel1_S | CCTCGGTGATAGGCACTTAAAC | RT-qPCR | This study |
| PIR-GM-Tel1_AS | AAAGGGACCAACTCTGGAATG |  |  |
| PIR-GM-PGK1-S | ATGTCTTTATCTTCAAAGTTGT | RT-qPCR | Garg et al, 2019 |
| PIR-GM-PGK1-AS | GGTTGGCAAAGCAGC |  |  |
